# Instinct to insight: Neural correlates of ethological strategy learning

**DOI:** 10.1101/2023.09.11.557240

**Authors:** Kai Lu, Kelvin T. Wong, Lin N. Zhou, Yike T. Shi, Chengcheng J. Yang, Robert C. Liu

**Author notes:** Corresponding author: Robert C. Liu.

## Abstract

In ethological behaviors like parenting, animals innately follow stereotyped patterns of choices to decide between uncertain outcomes but can learn to modify their strategies to incorporate new information. For example, female mice in a T-maze instinctively use spatial-memory to search for pups where they last found them but can learn more efficient strategies employing pup-associated acoustic cues. We uncovered neural correlates for transitioning between these innate and learned strategies. Auditory cortex (ACx) was required during learning. ACx firing at the nest increased with learning and correlated with subsequent search speed but not outcome. Surprisingly, ACx *suppression* rather than facilitation during search was more prognostic of correct sound-cued outcomes – even before adopting a sound-cued strategy. Meanwhile medial prefrontal cortex encoded the last pup location, but this decayed as the spatial-memory strategy declined. Our results suggest a neural competition between a weakening spatial-memory and strengthening sound-cued neural representation to mediate strategy switches.

## Introduction

Ethological behaviors operate instinctively, with animals predisposed towards innate strategies for deciding on responses to conspecific signals or threats^1,2^, though “open genetic programs”^3^ allow experience to adaptively alter strategies. Despite progress understanding goal-directed decision-making^4^, including switching between established strategies^5^, how the mammalian brain changes while learning from scratch to displace an innate strategy with a more efficient one in behaviors sculpted by evolution remains unexplored. We model ethological strategy-learning in a reproductive behavior. Mice make choices while searching for pups to retrieve. Retrieval is a hard-wired motor program that can be optogenetically activated – even when latent in pup-naïve males and virgins^6,7^. Yet *how* animals search involves strategic decisions influenced by experience. An innate strategy when few directional cues exist is to return to where a pup was last found^8–10^ – a spatial-memory form of the “win-stay/lose-switch” strategy^11,12^. However, mice can learn to use sounds for searching^13,14^, though how neural activity supports successful search is unknown. We used electrophysiology in freely moving, pup-retrieving mice to discover how plasticity within auditory cortex (ACx) and the medial prefrontal cortex (mPFC), which regulates spatial working memory^15^ and pup retrieval in rodents^16,17^, relates to the learned adoption of a sound-cued versus spatial-memory strategy for pup search.

## Results

### Mice required ACx to learn to switch from an innate spatial-memory to a sound-cued search strategy

We trained virgin female mice over eight days to use a novel artificial sound to guide their search for pups in a T-maze^8,18^. Using a non-ethological sound ensured that we isolated how a *learned strategy* displaces an innate one without the confound that species-specific vocalizations might have predisposed modes of privileged sensory processing^19,20^.

Each trial started with a mouse having two pups in its nest at the T-maze base (Figure 1a). A bandpass (30-50kHz) amplitude-modulated (5 Hz) gaussian noise presented continuously from a speaker at an arm’s end (pseudorandomly chosen) indicated where a pup (taken from the nest) would be manually delivered after the mouse entered that arm. Playback then stopped, and mice invariably retrieved the pup back to the nest, providing a natural reset for the next trial as animals carried out their ethological behavior. Animals on Day 1 of training followed an innate spatial-memory strategy during the search phase of a trial, choosing the arm where they received the pup in the last trial 84±2% (mean ± standard error of the mean) of the time (Figure 1b). They then learned to use the sound cue to efficiently search for pups (Figure 1c) within 3-7 days (Figure 1d), during which the use of the initial spatial-memory strategy dropped to 45±1%, near chance.

**Figure 1.**
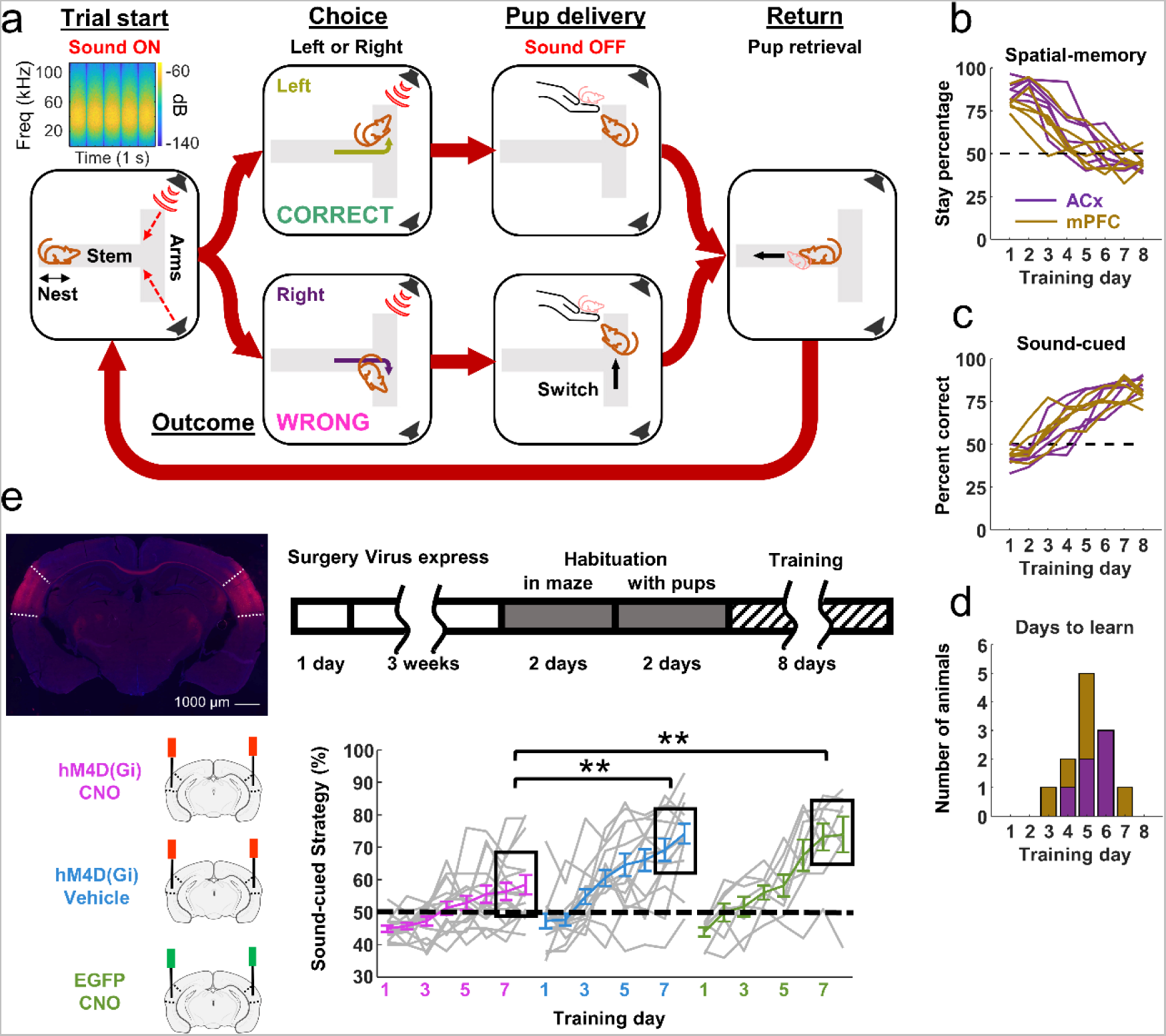
Animals required ACx to learn to switch from an innate spatial-memory strategy to the novel sound-cued strategy to perform pup retrieval. a. *The task structure:* Each trial started when the subject mouse was at the nest when an AM-modulated band-pass noise (Top left, spectrogram) began playing continuously from one of the two arms. The experimenter delivered a pup for retrieval once the mouse chose the CORRECT arm (Top branch), ending the sound. If the WRONG arm was chosen (Bottom branch), the mouse always switched to the other arm and received the pup. Retrieval back to the nest initiated the next trial. b. *Assessment of spatial-memory strategy:* Stay percentage (150 total), which measures how often animals first chose to return to the prior pup location. All 12 implanted animals (Purple: ACx; Brown: mPFC) stopped using the spatial-memory strategy over the 8 training days. c. *Assessment of sound-cued strategy:* Percent of trials in which animals first chose the sound-cued side. All 12 animals consistently learned the auditory cue within 8 days. d. *Latency to learn:* Stacked histogram of day by which an animal reached significantly higher sound-cued success (binomial test, p < 0.05}). The majority of animals learned by Day 5 (Purple: ACx; Brown: mPFC). e. *Chemogenetic inactivation of ACx impaired learning:* Example bilateral hM4D(Gi) virus expression in ACx (Top left). Experimental timeline (Top right). The sound-cued performance (Bottom) of individual animals (grey lines) and mean of each day (colored lines with standard error of the mean error bars) for the hM4D(Gi) + CNO group (Magenta), compared with those of the hM4D(Gi) + Saline (Blue) and EGFP + CNO (Green). In the last two days, the performance of the hM4D(Gi) + CNO group was significantly lower than the hM4D(Gi) + Saline group (Wilcoxon rank sum test, z = −2.84, *p* = 0.005) and the EGFP + CNO group (Wilcoxon rank sum test, z = −2.52, *p* = 0.012).

Learning to use the sound-cued strategy required ACx activity. We chemogenetically silenced ACx bilaterally in a cohort of animals before each training session by expressing hM4D(Gi) in ACx and injecting CNO shortly before each training session (Figure 1e). We compared this group to controls that either received the same virus in ACx with vehicle injections, or were given an EGFP virus with CNO. Relative to both controls, hM4D(Gi)+CNO animals were significantly impaired in learning the auditory strategy. Hence, acquiring this task depends on cortex, and we next turned to uncovering the neurophysiological mechanisms within ACx and mPFC that promote a switch to a sound-cued search strategy.

### Overall average ACx and mPFC firing rates were stable over training

We recorded from twelve virgin female mice implanted with silicon probes in the left ACx (N = 6, Extended Data Figure S1a) or left mPFC (N = 6, Extended Data Figure S1b). After recovery from surgery and an additional day of habituation to the T-maze and pups (Figure 2a, bottom), recordings began on Day 1 of sound training with 150 trials/day. We recorded 1307 total units in ACx, of which 1279 units (829 single-units and 450 multi-units) were responsive to sounds (see Methods); we excluded sound non-responsive ACx units in all subsequent analyses. The overall average ACx spike rates when the sound was ON (Extended Data Figure S1c) or OFF (Extended Data Figure S1d) did not change significantly across training sessions. Furthermore, 966 units (519 single-units and 447 multi-units) recorded from mPFC also showed no significant differences in overall average spike rates across sessions when the sound was ON (Extended Data Figure S1e), or OFF (Extended Data Figure S1f). Hence, for both recording groups, the overall recording quality remained stable across training.

**Figure 2.**
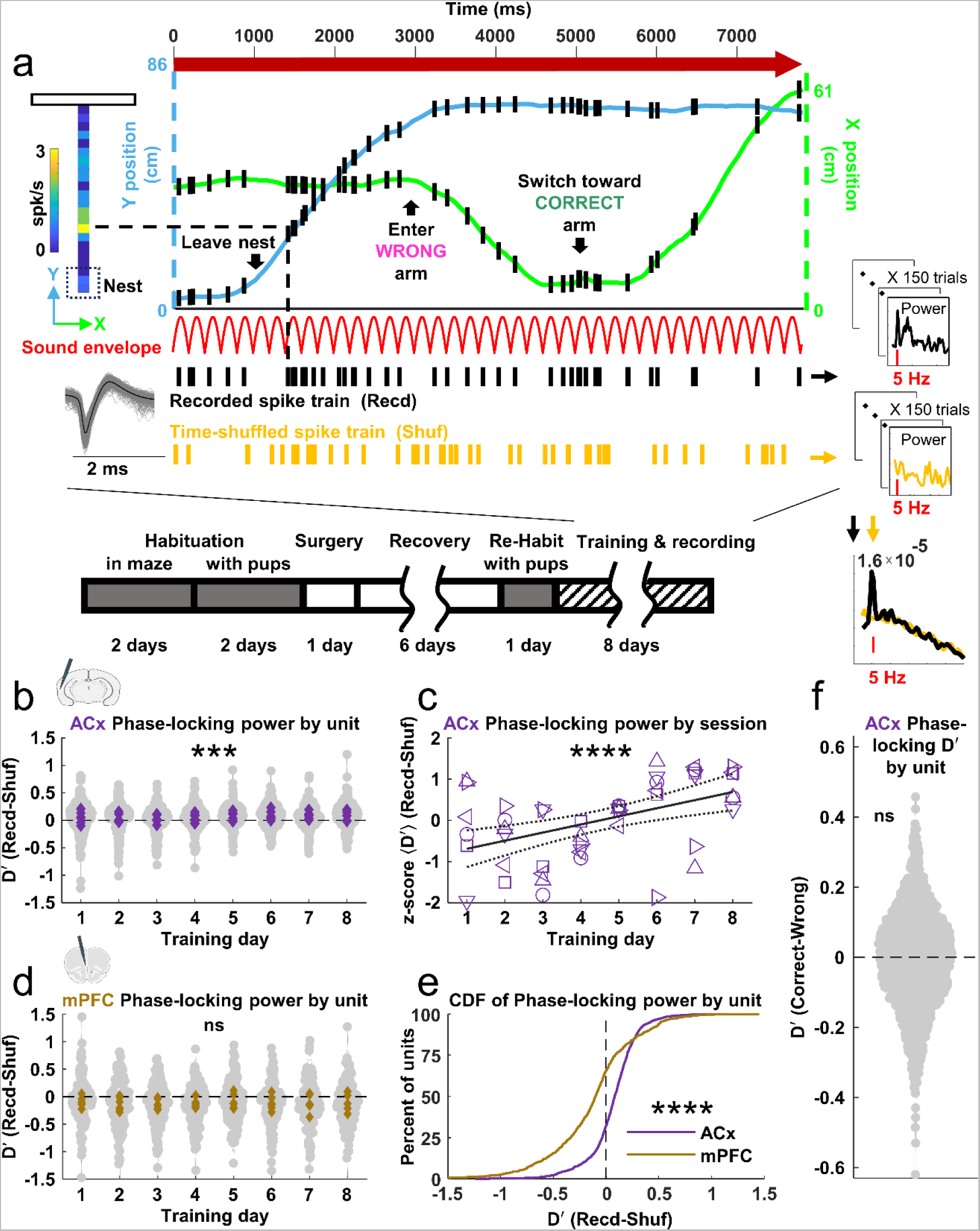
ACx, but not mPFC, units increased their 5 Hz phase-locking power in spike trains with training, without biasing outcome. a. *Single trial ACx firing during sound playback:* An ACx-implanted animal was recorded from Day 1 of training (Bottom, experimental timeline). Spiking (black hashes) of a single ACx unit (Middle left, spike waveform of all spikes) is illustrated as a function of the animal’s X (green) and Y (blue) position in the T-maze while the sound (envelope, red) played, with advancing time running from left to right. Spiking within a spatial bin was mapped onto the stem (Top left) to create spike rate heat maps for each trial. 5 Hz phase-locking power (Top right inset) of the spiking was calculated for each trial and compared (Bottom right inset) to the power (Middle right inset) after shuffling spikes in time (gold hashes). b. *ACx population’s 5 Hz phase-locking power across days by unit:* D’ between Recorded and Shuffled spike trains mildly increased during the 8 days training, tested across all units (linear mixed-effects model LMM, likelihood ratio LR= 9.396, *p* = 0.002). Here and in all similar figure panels (unless noted otherwise), each grey dot is a unit, each purple diamond indicates the mean across ACx units in a recording session, and asterisks show significance for changes across Days (*, *p* < 0.05; **, *p* < 0.01; ***, *p* < 0.005; ****, *p* < 0.001). c. *ACx population’s 5 Hz phase-locking power across days by recording session:* D’ between Recorded and Shuffled trials, averaged for all units recorded in a given session, were represented as within-animal daily means that were z-scored across the animal’s 8 days of training. Here and in all similar figure panels, each symbol type represents an individual animal. The linear regression fit with the 95% confidence interval (here and in all similar figure panels, black solid line flanked by black dotted lines) showed a significant increase in D’ across training days (linear regression, β = 0.197, *p* < 0.001). d. *mPFC population’s 5 Hz phase-locking power across days by unit:* D’ between Recorded and Shuffled spike trains showed no significant change across the 8 days training, whether by unit (LMM, LR = 0.113, *p* = 0.737) or by sessions (linear regression, β = 0.021, *p* = 0.723, data not shown). Here and in all similar figure panels, each brown diamond indicates the mean across mPFC units in a recording session. e. *Cumulative distribution of 5 Hz phase-locking power:* ACx and mPFC units differed significantly in the prevalence of phase-locking (hierarchical bootstrap, *p* <0.001). f. *Overall ACx phase-locking power did not bias outcome:* D’ between Correct and Wrong trials was not significantly different from zero (hierarchical bootstrap, *p* = 0.203) and did not change significantly across days (LMM, LR = 1.710, *p* = 0.191, data not shown).

### Phase-locking power in ACx but not mPFC increased over training yet did not systematically influence the sound-cued search strategy

During sound ON periods, beginning at the nest and ending upon pup delivery, ACx spiking occurred in both a temporally and spatially specific fashion along the T-maze stem. Temporally, units such as the one illustrated for Day 1 (Figure 2a) preferentially fired in-phase with the 5 Hz sound envelope. Overall, 38% of ACx sound-responsive units across all Days and animals showed significantly higher sound ON phase-locking power across trials when compared to the 5 Hz power when spikes were temporally shuffled. Phase-locking power increased modestly but significantly with training at the level of individual units (Figure 2b) and recording sessions (Figure 2c), suggesting a mildly improved ability for ACx to entrain to the sound. In contrast, only 20% of mPFC units showed significant phase-locking. Phase-locking power among mPFC units (Figure 2d) or sessions did not change over training, and was significantly less than in ACx (Figure 2e).

Despite ACx phase-locking changes, this neural activity measure was not systematically responsible for correctly using the sound to search on a trial-by-trial basis. On any trial, areas downstream of ACx may simply act on a stable (albeit potentially noisy) ACx representation of the sound to decide whether to follow it in choosing an arm. Alternatively, variable ACx activity itself could directly bias the use of the sound in searching if that activity systematically differs between subsequent Correct versus Wrong sound-cued trial outcomes. To assay this for phase-locking, we computed a D’, which normalizes mean differences by variance across trials^21^, between Correct and Wrong trials for each unit across all days and found that D’ was not consistently different than 0 (Figure 2f) and did not change with training. Hence, better ACx phase-locking with learning improved the sound’s temporal coding but did not systematically bias the use of a sound-cued strategy for pup search.

### Relative sound-evoked nest firing increased over training for ACx but not mPFC units yet did not systematically influence the sound-cued search strategy

Spatially, both ACx and mPFC units tended to fire preferentially in particular locations along the stem. We constructed a location-dependent, z-scored firing rate map for each trial (Figure 2a) for each unit. On Day 1, ACx units tended to have their highest firing rates close to the intersection of the T-maze stem with the arms (Figure 3a, marked “T”), which is where both speakers were pointed. By Day 8, many units had acquired higher relative firing rates at the nest (bins 1-3, marked “N”), even though this location was further from the speaker. The average relative nest firing rate significantly increased during training, tested both at the level of all individual units (Figure 3b) and all recording sessions (Figure 3c). This was irrespective of unit best frequency (BF, Spearman correlation, rs = −0.073, *p* = 0.108). In contrast, maximum mPFC firing rates when the sound was ON were located close to the nest on Day 1 (Figure 3d), and remained there even on Day 8. Across all mPFC units there was no modulation of relative nest firing by training day (Figure 3e), even though these units showed sustained high nest responses across all days. Though it increased with training, the ACx’s relative nest firing was not consistently responsible for correctly using the sound as a search strategy on a trial-by-trial basis, as inferred from the D’ centered near zero (Figure 3f).

**Figure 3.**
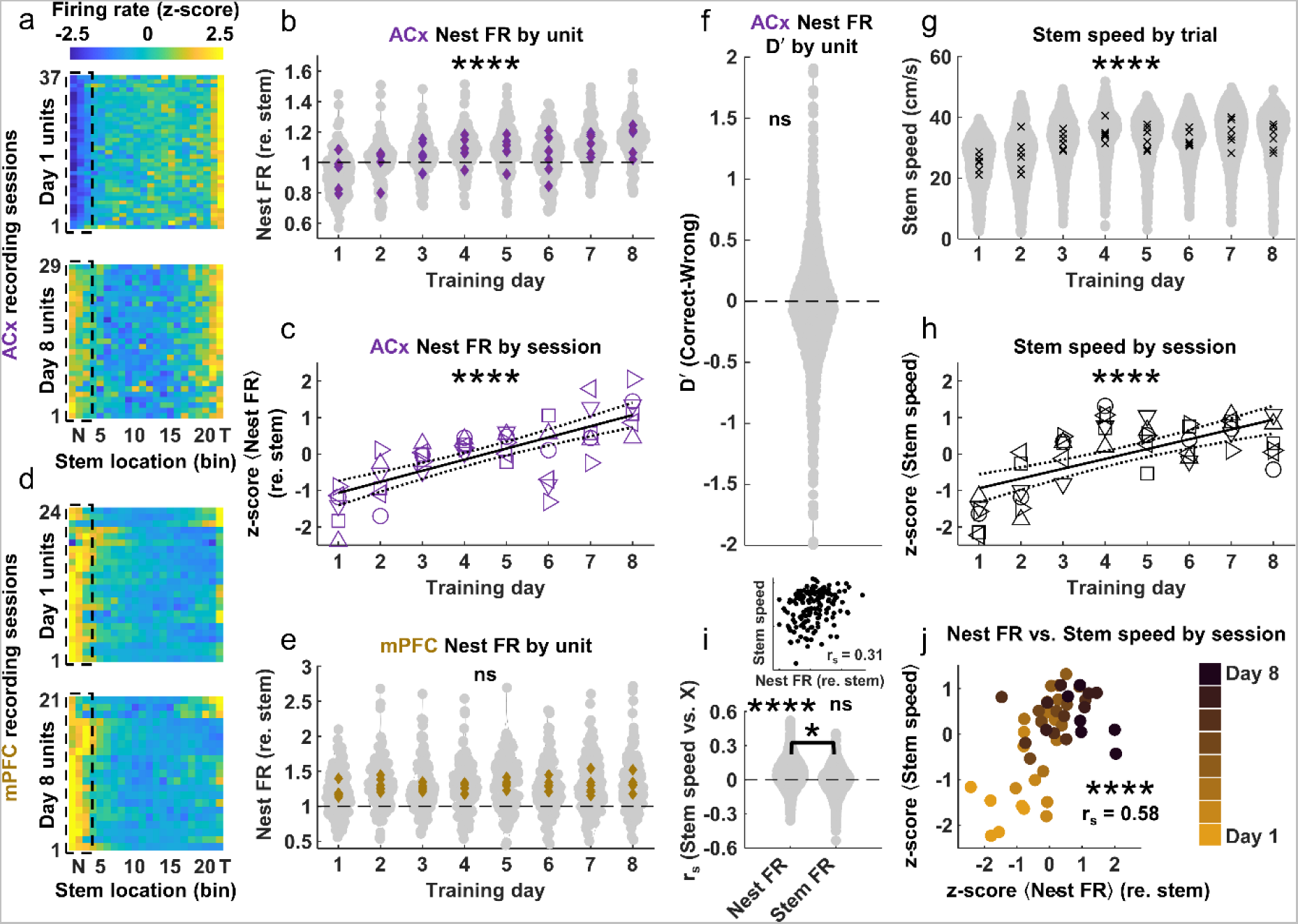
Relative nest firing in ACx but not mPFC increased over training and was correlated with increased search speed. a. *Location-dependent firing of simultaneously recorded ACx units:* Sound ON, trial-averaged firing rate heat map as a function of stem location (including the nest but not arms) for all recorded ACx neurons during a Day 1 (Top) and a Day 8 (Bottom) training session. To visualize the spatial pattern of different units in the map, spike rates were z-scored across all spatial bins for each unit. Nest responses (dashed black box) were higher on Day 8 than in Day 1 in this animal. b. *ACx population’s relative nest firing rate across days by unit:* Firing rate at the nest, normalized within-unit to its average firing rate in the stem, increased significantly across days (LMM, LR = 246.584, *p* < 0.001). c. *ACx population’s relative nest firing rate across days by recording session:* Relative firing rate at the nest, averaged for all units recorded in a session and represented as within-animal daily means that were z-scored across the animal’s 8 days of training, significantly increased across days (linear regression, β = 0.305, *p* < 0.001). d. *Location-dependent firing of simultaneously recorded mPFC units:* Same as in (a) for mPFC units from an example animal. e. *mPFC population’s relative nest firing rate across days by unit:* mPFC units sustained higher Sound ON firing rates in the nest compared to in the stem (hierarchical bootstrap, *p* < 0.001 for all days), but did not change across days (LMM, LR = 1.802, *p* = 0.180). f. *Overall ACx relative nest firing did not bias outcome:* D’ between Correct and Wrong trials was not significantly different from zero (hierarchical bootstrap, *p* =0.1521). g. *Animal’s stem speed across days by trials:* Locomotion speed as animals traversed the stem significantly increased across days (LMM, LR = 998.285, *p* < 0.001). Each grey dot represents a trial and each black X indicates the average across trials in a recording session. h. *Animal’s stem speed across days by recording session:* Animals ran down the stem during the Sound ON search phase significantly faster as training progressed over the 8 days (linear regression, β = 0.269, *p* < 0.001). Trial-averaged daily means were z-scored across days within each animal. i. *Correlation between speed and relative nest firing rate at the unit level:* Example of correlation between each trial’s stem speed and nest firing rate (FR) for an example unit (Top inset). Correlation coefficients between stem speed and a unit’s nest firing rate (Left violin) was significantly greater than zero (hierarchical bootstrap, *p* < 0.001) and higher than the correlation with the firing rate along the stem itself (Right violin, hierarchical bootstrap, *p* < 0.001). The latter was not significantly different from zero (hierarchical bootstrap, *p* = 0.383). j. *Correlation between speed and relative nest firing rate at the recording session level:* Scatter plot of z-scored unit-averaged relative nest firing rate from a session as a function of the z-scored trial-averaged stem speeds shows a significant positive correlation (Spearman correlation, r_s_ = 0.58, *p* < 0.001). Here and in all similar figure panels, each dot represents a recording session and the color of each dot indicates different training days.

### Search speed increased over training and specifically correlated trial-by-trial with stronger relative nest ACx firing

Despite the lack of obvious impact on search strategy, the relative ACx nest firing nevertheless revealed how subjects carried out trials. How quickly animals ran down the stem (bins 4-22) after exiting the nest systematically increased over training either by trial (Figure 3g) or session (Figure 3h), reaching ∼34 cm/s. Importantly, the average correlation between the trial-by-trial speed *along the stem* and a unit’s trial-by-trial firing rate *in the nest* (Figure 3i, inset) was significantly higher than zero (Figure 3i). In fact, the ability to explain each trial’s stem speed by using the nest firing was better than by the firing rate *along the stem* itself (*r_s_*(stem speed vs. nest firing rate) > *r_s_*(stem speed vs. stem firing rate), the latter which was ∼0). Finally, average stem speed (z-scored across days within animal) and average population nest response (z-scored across days within animal) were significantly correlated over training (Figure 3j). Hence, as animals learned the sound’s behavioral relevance, an overall increased ACx response in the nest may have helped to detect the sound farther from the speaker, enhancing arousal and prompting more vigorous, rapid pup search.

### Stem firing rate in ACx but not mPFC was prognostic of correct outcomes, systematically influencing the sound-cued search strategy

Since neither ACx phase-locking nor relative nest firing consistently influenced the adoption of a sound-cued rather than a spatial-memory search strategy for pups, we turned to the *stem* firing rate once animals exited the nest. An individual ACx unit’s spatially binned firing rate could be bidirectionally predicted by multiple factors, including whether sounds played from the Left versus Right, but also whether the trial’s outcome was Correct versus Wrong (Figure 4a). To rigorously quantify such varied forms of modulation, we fit each unit’s spatially binned, trial-by-trial stem firing rate to a Generalized Linear Model (GLM)^22^ with location, sound source side and level (roved over a 4 dB suprathreshold range) and outcome as predictors (Figure 4b). Using a model comparison approach (see Methods), we found that virtually all (96%±1) ACx units per recording session were significantly modulated by location, while 44±2% were by sound side and 39±2% by sound level (Figure 4c) – as might be expected for acoustically responsive neurons.

**Figure 4.**
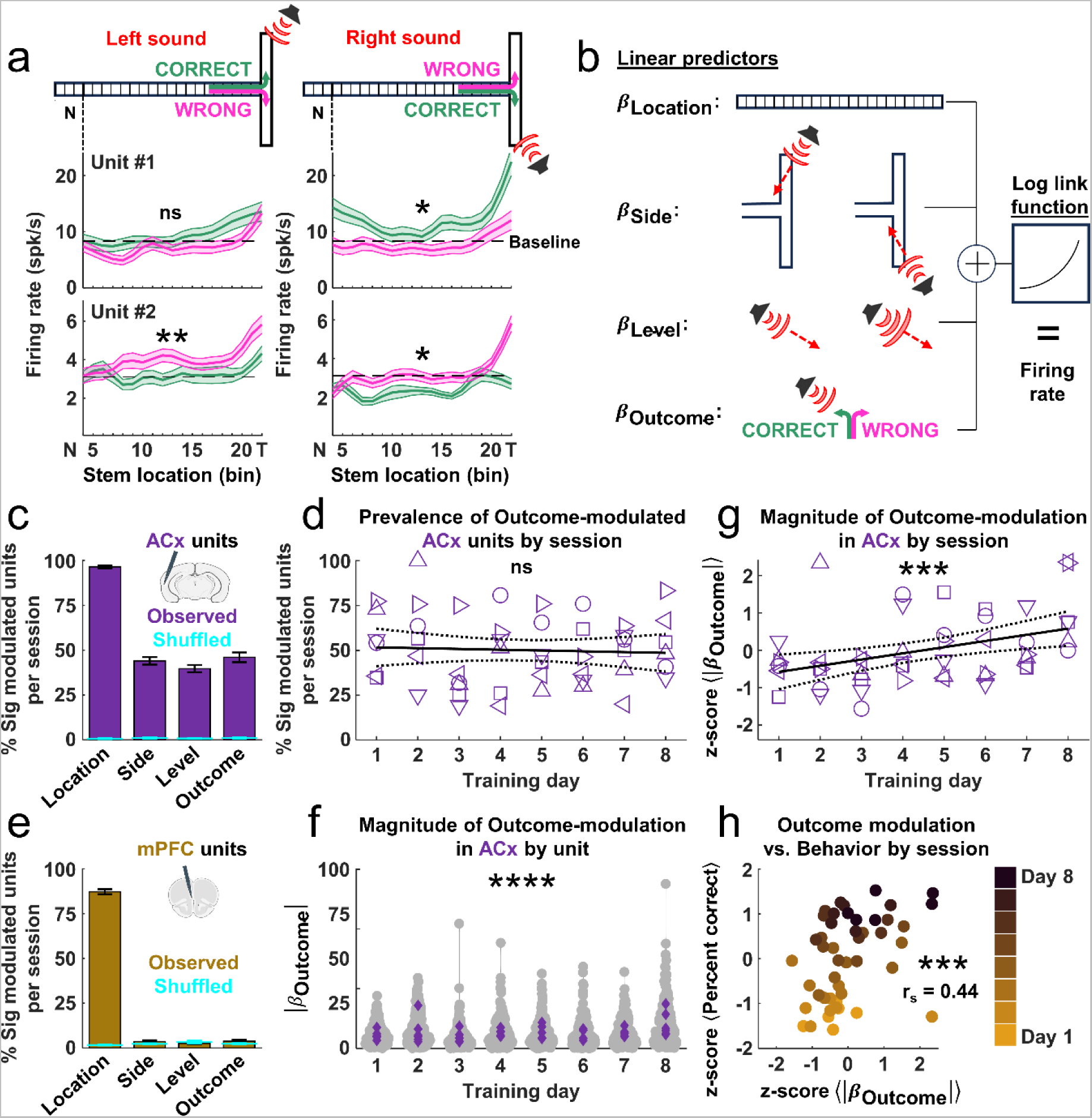
Stem firing in ACx was increasingly modulated by the sound-cued Outcome. a. *Modulation of ACx firing rate by multiple factors:* Firing rate of two ACx units (Middle and Bottom row), broken down and averaged by position along the stem (Top) depending on whether the Sound is presented from the Left (Left column) or Right (Right column) and the animal subsequently had the Correct (Green) or Wrong (Pink) Outcome. Average baseline firing rate during Sound OFF phase shown as a black dashed line. b. *Linear predictors for generalized linear model of unit firing rate:* Four factors – location, the sound side and level, and trial outcomes – were used to predict the spike rate in each spatial bin along the stem, with a log link function. c. *Prevalence of significantly modulated ACx units:* Percent of units observed in each recording session showing significant modulation by each factor, as modeled by GLMs (likelihood ratio test, *p* < 0.05/(number of locations=22)). Error bars reflect standard error of the mean across recording sessions. All factors modulated a larger proportion of ACx population than if trials were randomly shuffled (Cyan). d. *Prevalence of significantly modulated ACx units across days by recording session:* The percent of units in ACx in each training session that showed significant effect of outcomes was above 50% even on the first day and did not significantly change across days (linear regression, β = −0.004, *p* = 0.723). e. *Prevalence of significantly modulated mPFC units:* As in (c) for mPFC units. While mPFC firing clearly depended on location for nearly all units, sound side, level and outcome did not modulate the mPFC population at levels different from when trials were shuffled (Wilcoxon signed-rank test: *p* > 0.05, for all three factors). f. *Magnitude of outcome-modulation of ACx across days by unit:* The strength (absolute value) of the GLM coefficient β for outcome-modulation significantly increased over days (LMM, LR = 19.808, *p* < 0.001). g. *Magnitude of outcome-modulation of ACx across days by recording session:* The strength of the GLM β coefficients for outcome-modulation, averaged for all units recorded in a session and represented as within-animal daily means that were z-scored across the animal’s 8 days of training, significantly increased across days (linear regression, β = 0.166, *p* = 0.004). h. *Correlation between behavioral outcome and neural outcome-modulation at the recording session level:* Scatter plot of z-scored unit-averaged GLM β_Outcome_ coefficients from a session as a function of the z-scored daily behavioral performance using the sound-cued strategy showed a significant positive correlation (Spearman correlation, r_s_ = 0.445, *p* = 0.002).

Intriguingly, on average 46±3% of units were significantly predicted by the *subsequent* “outcome” of the trial and were thus “prognostic” of whether the animal would correctly use the sound in their decision strategy. Such prognostic units were present at high levels on Day 1 (Figure 4d) and the proportion remained high throughout training, implying the unexpected finding that ACx activity discriminated Correct outcomes even before animals behaviorally demonstrated ∼Day 3-7 that they had learned to use the sound (Figure 1d). For comparison, in mPFC (Figure 4e), while 87±1% of units showed a significant effect of spatial location, few (<4%) showed significant modulation by sound side, level, or outcome while the sound was ON, and the proportions were not different from trial-shuffled estimates.

Although the prevalence of prognostic ACx units was stable across days, the modulation strength varied over training. There was a significant effect of training day on the absolute value of the coefficients for the GLM outcome factor across the neural population (Figure 4f). Fitting the means from each recording session, the magnitude of the outcome factor increased significantly with training (Figure 4g), indicating that individual ACx stem firing became progressively more indicative of whether the animal would approach the sound. Across sessions, prognostic modulation improved along with daily performance in the T-maze. The magnitude of the outcome modulation averaged in each recording session was positively correlated with the performance across all recording sessions (Figure 4h). Thus, the GLM analysis discovered that stem firing in ACx but not in mPFC was prognostic of the up-coming sound-cued outcome, both within and across training sessions, from the earliest days of training, with a magnitude that correlated with animals’ performance across days.

### Overall modulation of ACx stem firing by sound-cued outcome was suppressive and not related to other factors

Using a GLM entails assumptions as to how predictors impact the firing rate, so to assess the model-free effect of outcome on ACx firing, we calculated the D’ for the stem-averaged firing rate between Correct and Wrong trials for all units. Surprisingly, although Correct outcomes could either enhance or suppress the firing rate, on nearly every training day (Figure 5a) more units showed *lower* spike rates for Correct trials. Hence, 72% of units had D’ values that were negative, so that the overall median D’ was significantly less than zero (Figure 5b). On the other hand, D’ for side choice was symmetric around zero. Importantly, an overall suppressive effect on outcome D’ appeared even in the first training day (Day 1 in Figure 5a) *before* animals learned the sound-pup association, consistent with the GLM results.

**Figure 5.**
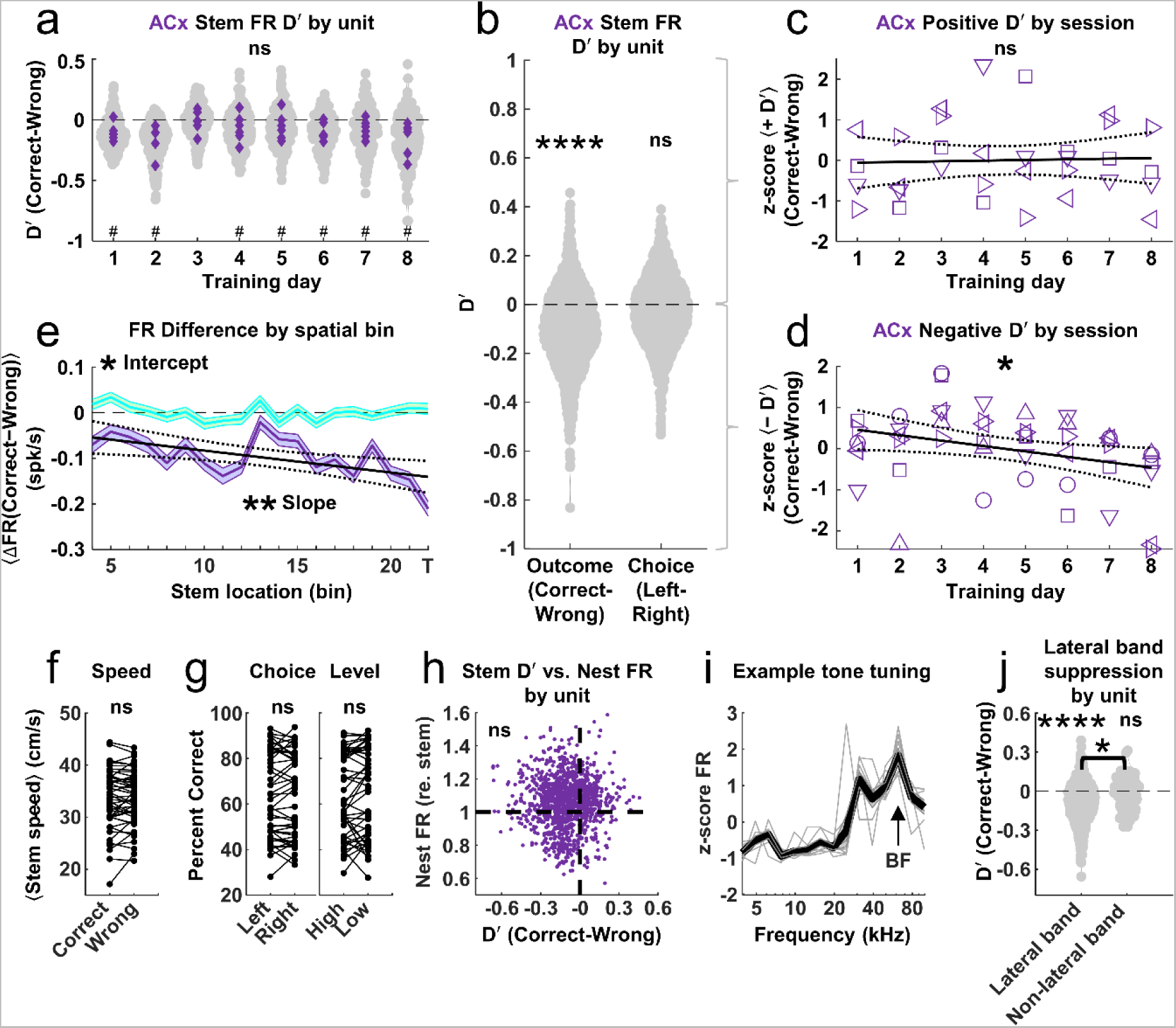
Overall modulation of ACx stem firing by sound-cued outcome was suppressive and stronger in lateral band units. a. *ACx outcome-modulation across days:* D’ between firing rates from Correct and Wrong trials were significantly lower than zero even on Day 1 (hierarchical bootstrap, *p* < 0.001), and did not change significantly across days, tested either by unit (LMM, LR = 0.042, *p* = 0.838) or by recording session (linear regression, β = −0.043, *p* = 0.475, data not shown). #, *p* < 0.05 for hierarchical bootstrap test of each day. b. *Overall outcome and choice modulation of ACx firing:* D’ between ACx firing rates from Correct and Wrong trials showed an overall suppressive effect of Correct Outcomes (Left, hierarchical bootstrap, *p* < 0.001). D’ between ACx firing rates from Left and Right choices was not significantly different from zero (hierarchical bootstrap, *p* = 0.163). c. *ACx units positively modulated by outcome across days:* Units whose average firing rate was larger for Correct versus Wrong trials, as inferred from a positive D’ for outcome, remained unchanged across days, tested either by unit (LMM, LR = 2.850, *p* = 0.091, data not shown) or by recording session (linear regression, β = 0.016, *p* = 0.832), where the ACx units with positive D’ in each session were averaged and z-scored across days within each animal. d. *ACx units negatively modulated by outcome across days:* Units whose average firing rate was larger for Wrong versus Correct trials, as inferred from a negative D’ for outcome, became more strongly modulated across days, tested either by unit (LMM, LR = 9.700, *p* = 0.002) or by recording sessions (linear regression, β = −0.132, *p* = 0.025). e. *ACx outcome-modulation across stem location:* Population means of spike rate differences (Correct - Wrong) across spatial bins along the stem (Purple), compared with those obtained from shuffled trials (Cyan). Significant suppression by Correct outcomes appeared in all bins along the stem (linear regression, Intercept = −0.049, *p* = 0.014), with greater suppression as animals moved towards the arms (linear regression, β = −0.005, *p* = 0.008). f. *Outcome effects on stem speed by session:* Average locomotion speeds along the stem for Correct and Wrong trials were not significantly different (Wilcoxon signed-rank test, z = 0.390, *p* = 0.697). g. *Choice and level effects on sound-cued outcome by session:* The proportion of Correct trials in a session did not systematically depend on side choice (Left versus Right, Wilcoxon signed-rank test, z = 0.010, *p* = 0.992) or sound level (High versus Low volumes, Wilcoxon signed-rank test, z = −1.513, *p* = 0.130). h. *Correlation between relative nest firing rate and outcome-modulation at the unit level:* D’ between firing rates from Correct and Wrong trials was not correlated with the relative nest firing rate across all ACx units (Spearman correlation, rs = 0.014, *p* = 0.608). i. *Passive-listening tuning curve:* Example pure tone tuning curve obtained from a muti-unit recording channel in one animal across all 8 days (Grey lines). The black trace shows the mean across all days. The channel’s BF extracted from the average tuning curve is indicated by the arrow. j. *Stronger outcome suppression in lateral band units:* D’ between firing rates from Correct and Wrong trials was significantly below zero for units tuned to BFs lateral to the Sound’s spectrum (hierarchical bootstrap, *p* < 0.001), and significantly more negative than those from non-lateral band units (hierarchical bootstrap, *p* < 0.048, Wilcoxon rank-sum test: z = −3.241, *p* = 0.001). D’ for non-lateral band units was not significantly different from zero (hierarchical bootstrap, *p* = 0.093).

Across days, the overall D’ did not change significantly. However, since we observed an increase with training in the absolute magnitude of outcome modulation as derived from the GLM, we split the units according to whether their D’ was enhanced (positive) or suppressed (negative) for Correct outcomes to determine whether the magnitude of these subpopulations changed. For units with positive D’, we could not detect a significant difference across days either on a per unit or per session (Figure 5c) basis. For negative D’ units though, D’ became progressively more negative later in training, both on a per unit and per session basis (Figure 5d). Hence, training was particularly effective in increasing the degree of prognostic suppression of the ACx stem-averaged firing rate for Correct relative to Wrong trials.

When broken down by spatial location along the stem, the overall suppression by Correct outcome was apparent from the start of the stem and became more negative as animals approached the T-maze arms (Figure 5e). Although ACx suppression has been linked to engagement and movement^23–25^, animals in our paradigm were always engaged in the retrieval task and moved with statistically similar speeds on both Correct and Wrong trials in a session (Figure 5f). Moreover, while some individual ACx units (Figure 3i, inset) showed a significant (*p* < 0.05) Spearman correlation between speed and firing rate, similar proportions were positive (6%) and negative (8%) so that this was unlikely to explain our dominant prognostic suppression. Side choice and sound level (Figure 5g) were also not systematically related to achieving a Correct outcome.

Finally, since the relative nest firing rate also changed across days, we investigated whether it could systematically explain prognostic outcome modulation of the stem firing rate. Across the population of all units, we found no correlation between these measures (Figure 5h). Along with the fact that D’ for outcome modulation of the relative nest firing itself was not consistently different than 0 (Figure 3f), our data indicate that ACx firing *at the nest*, while evidently related to vigorous pup search, does not systematically reveal the strategy of using the sound as a basis for searching.

### Prognostic suppression of ACx stem firing was stronger in lateral band units

Having a Correct sound-cued outcome be associated with *less* ACx activity along the stem compared to Wrong outcomes is contrary to the usual expectations for the role that ACx plays in mediating sound perception^26^. One potential explanation is that Correct outcomes are mainly associated with suppression of neurons attuned to frequencies outside the sound’s spectrum. In fact, earlier work in passive listening animals has shown that behaviorally salient natural ultrasonic vocalizations evoke strong “lateral band suppression” of neurons with BFs just below the ultrasonic frequency range^27^. To investigate this possibility, for four ACx animals we included a passive listening condition before each training session when we played pure tones to estimate a channel’s consistent overall BF (Figure 5i). For the population of ACx units falling into the lateral band (BFs were 0.3 to 2 oct lateral to the 40 kHz central frequency of the sound), the average D’ was significantly below zero (Fig 5j, left) and was significantly less than the D’ for units not in the lateral band, whose average D’ was not different from zero (Fig 5j, right). Hence, prognostic suppression of lateral band firing along the stem may help increase the signal-to-noise in the ACx population neural representation of the sound by reducing the activity of neurons whose BFs do not reflect its spectral content, allowing ACx to have a systematic impact on the search strategy.

### The firing rate in a subset of mPFC units retained a side preference during return

Our results thus far suggest that changes over training in the prognostic suppression of lateral band BF neurons in ACx while animals ran down the stem, but not ACx firing in the nest, contributed to adopting the novel sound-cued search strategy. Yet if a mechanism that correlates with successful use of an auditory strategy is present at high levels already on Day 1, why does it still take 3-7 days for animals to behaviorally switch their strategy? One possibility is that the weight of this mechanism in biasing downstream decisions for releasing behavior must still build up, which we observed, but another is that neural correlates of the initial spatial-memory strategy dominate in these early days and need to weaken before the more efficient strategy can take hold. We thus turned next to investigating how mPFC activity, which has been found to feature social-place fields^28^, encodes the memory of the location it received the pup on the last trial and changes over training.

During a trial, once a pup was delivered for retrieval, the sound was turned OFF and the animal ran back through the arm and stem to deposit the pup at the nest. Some units began firing in the arm and continued spiking, albeit usually with a declining rate, as the animal traversed the path back along the stem (Figure 6a, time increasing right to left). The firing rate could systematically differ depending on whether the animal started in the Left or Right arm, as seen for two example units (Figure 6b), and that preference could be maintained during the return along the stem (Figure 6c). Among any given recording session’s units, there was typically a continuum of firing rate differences between starting from the Left versus Right (Figure 6c-d), with 33% of all mPFC units showing a significant sustained preference for one arm over the other (Wilcoxon rank-sum test, *p* < 0.05, tested per unit). Even when we removed the 18% of all mPFC units that showed potential premotor activity related to preferential turning (see Methods), 27% of the remaining units still showed a significant sustained preference for one arm or the other. Importantly, such units were present on Day 1, where this arm selectivity could have supported a neural substrate for a working memory^29^ of the last rewarded side.

**Figure 6.**
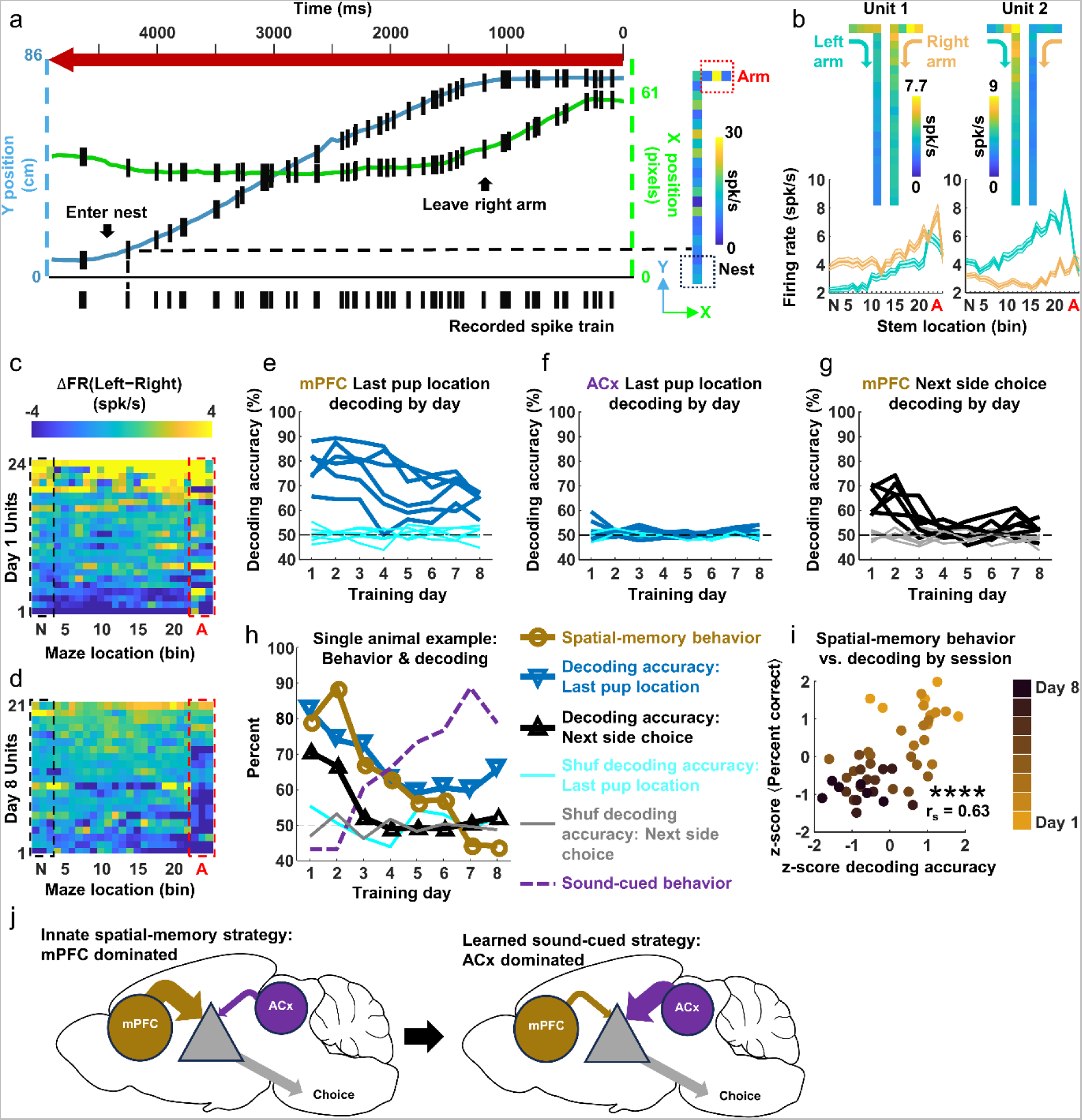
Firing rates during retrieval for a subset of mPFC units initially decoded the last pup and next choice sides but lost that as the spatial-memory strategy declined. a. *Single trial mPFC firing during pup retrieval:* A mPFC-implanted animal was recorded from Day 1 of training. Spiking (black hashes) of a single mPFC unit is illustrated as a function of the animal’s X (green) and Y (blue) position in the T-maze as the animal returned to the nest, with advancing time running from right to left. Spiking within a spatial bin was mapped onto the stem and T-maze arm (Right) to create spike rate heat maps for each trial. b. *mPFC units fire differentially when animals return from different arms:* Firing rate heat maps (Top) and spatial location dependence (Bottom) for two mPFC units (Left and Right) during return trip to nest while retrieving pups. Trials for returning from the left or right side were averaged separately. Unit 1 showed sustained higher spike rates when retrieving pups from the right arm. Unit 2 showed the opposite. c. *Arm preference across mPFC population recorded on Day 1:* Heat maps of spike rate differences between left and right arm-returning trials in Day 1 of an example animal. d. *Arm preference across mPFC population recorded on Day 8:* Same as (c) except for Day 8 of the same animal. e. *mPFC neural population decoding accuracy of the last pup location by day:* On Day 1, decoding accuracy (based on firing rates during the silent return to the nest) across the six mPFC-implanted animals (Dark blue) was around 80%, and accuracy reduced to ∼60% by Day 8, but remained significantly higher than decoding accuracy from shuffled (Cyan) trials (ANOVA, main effect of observed/shuffled, F(1,40) = 36.015, *p* = 0.002; interaction between observed/shuffled and days: F (7,40) = 10.446, *p* < 0.001). f. *ACx neural population decoding accuracy of the last pup location by day:* Decoding accuracy of the last pup location by ACx units was not different from shuffled (ANOVA, main effect of observed/shuffled, F(1,40) = 3.251, *p* = 0.131; interaction between observed/shuffled and days (F (7,40) = 0.929, *p* = 0.497). g. *mPFC neural population decoding accuracy of the next side choice by day:* On Day 1, decoding accuracy (based on firing rates during the silent return to the nest) across the six mPFC-implanted animals (Black) was significantly different from 50% chance (Wilcoxon signed-rank test, z = 4.985, *p* < 0.001), significantly different from shuffled (Grey) trials (ANOVA, F(1,40) = 68.820, *p* <0.001), and reduced significantly over days (ANOVA, interaction between observed/shuffled and days: F (7,40) = 6.310, *p* < 0.001). Decoding accuracy of the next side choice was also significantly lower that the decoding accuracy for the last pup location (Wilcoxon signed-rank test, z = 5.990, *p* < 0.001). h. *Single animal illustration of relationship between behavioral performance and mPFC neural decoding:* The decoding accuracy of last pup location, next side choices, spatial-memory behavior, and sound-cued behavior in a session changed over days in a correlated manner. i. *Correlation between spatial-memory behavioral outcome and neural decoding accuracy at the recording session level:* Scatter plot of z-scored decoding accuracy of the last pup location averaged from a session as a function of the z-scored daily behavioral performance using the spatial-memory strategy showed a significant positive correlation (Spearman correlation, rs = 0.627, *p* < 0.001). j. *Conceptual neural model of switching from an innate to a learned strategy.* Initially, mPFC (Gold oval) carries the weight for the innate spatial-memory strategy and drives behavioral choices while searching for a pup (Top). ACx (Purple circle) carries the weight of new sensory information (Bottom), which can bias the animal’s choice (as reached through a hypothesized downstream region, Grey triangle) even on Day 1 of training. An early, dominant role played by mPFC switches to ACx domination during learning.

### Population mPFC activity during return initially decoded last-trial and next-choice sides but declined with a loss of the spatial-memory strategy

We hypothesized that side selectivity in mPFC units helped encode at the population level spatial information about the last pup location in the just-concluded trial to influence the next side choice when animals searched for pups using a spatial-memory strategy. Indeed, population mPFC (Figure 6e, 78.4±8% mean decoding accuracy) but not ACx (Figure 6f) firing rates during retrieval decoded the prior pup location well on Day 1. Moreover, the decoding accuracy of the next choice (Figure 6g) was significantly above chance but lower than that for last pup location – consistent with an existing but reduced utility of the mPFC activity, during the return trip, to reveal the future decision during the following trial.

We next asked whether mPFC activity stably encoded the spatial-memory strategy across training, as might be expected if pup-associated auditory neural information simply had to become sufficiently amplified to overcome the innate strategy. Intriguingly though, mPFC decoding accuracies for both last pup location and next side choice declined after ∼Day 2, just as the usage of a spatial-memory strategy dropped, as illustrated for an example animal (Figure 6h). This brain-behavior correlation held across all recording sessions for all animals, with both measures declining over training (Figure 6i). Hence, mPFC activity retained a spatial mnemonic role in guiding search only as long as the animals utilized a spatial-memory strategy, consistent with the hypothesis that this role’s weakening helps facilitate the adoption of a new sound-cued search strategy.

## Discussion

Decisions are inherent in natural social interactions, but their underlying neural basis can be opaque due to their unconstrained nature. Here we took advantage of a highly stereotyped reproductive behavior in rodents – pup retrieval – to uncover neural correlates of learning a new strategy for making more efficient decisions in an ethological social context. As female mice learned that a novel sound was more reliably predictive of where a pup would be delivered than the location where a pup was last found, the ACx and mPFC were engaged during distinct behavioral stages and plastic across days to systematically reflect the animal’s decision to use the sound to guide search. ACx relative firing in the nest correlated with more vigorous searching, while, paradoxically, ACx *suppression* along the stem was prognostic of subsequent success in choosing the sound-paired location. Both neural correlates strengthened with experience, although prognostic suppression was unexpectedly prevalent already on Day 1 – before adopting a sound-cued strategy. Meanwhile, mPFC firing while the sound was ON did not contribute to Correct sound-cued decisions, but instead on Day 1 retained during the return trip an encoding of the last pup side – a representation of the animal’s innate spatial-memory strategy that weakened before the sound-cued strategy was adopted. These results illuminate how the neural correlates of competing search strategies evolve dynamically during a natural behavioral sequence and change over learning to support switching to a more efficient way of making decisions in a social behavior.

Despite a wealth of research into the neural bases for perception, decision-making and learning^4,30,31^, there is little prior work in mammals on neural mechanisms for *learning strategies* to make decisions in natural behaviors. Behavioral strategies are rules animals follow to choose between uncertain options to reach goals. Rather than being purely random, many ethological strategies in innate behaviors follow stereotyped patterns of choices^32,33^. A classic example is the win-stay/lose-switch strategy for bandit problems^34^, which encompass our T-maze paradigm. All our mice innately followed such a strategy by returning to the prior pup-rewarded arm on Day 1, and immediately switching to the other arm if no pup was given – even though this was not efficient given the pseudorandom trial structure. They learned to use the sound-based strategy for *choosing* an arm but did not have to learn any new behaviors to receive their reward. Hence, without having to instrumentally shape a new motor program, we observed how experience from the first moment of hearing a pup-associated sound changes the strategic framework for decision-making, and uncovered neural correlates of that change. This differs from prior studies of associative stimulus-action learning^35^, perceptual learning to improve stimulus detection and/or discrimination^26,36^, or task-switching^37–39^, since those involve first learning new motor actions, thus occluding investigation of how the neural substrates for innate decision strategies in ethological behaviors are altered by experience.

Amongst ethological reproductive behaviors in rodents, pup tactile and chemosensory cues act as innate key stimuli to control pup-directed responses in naïve virgins^40^, while pup sounds become associated to pups through pup care experience to release maternal care, even in co-caring virgins^13^. Pup care experience correlates with plasticity in ACx’s response to the natural pup sounds^14,27,41–44^, and these changes may help regulate maternal motivation to approach pups^45^. Our results support this view in that increased ACx relative nest firing predicted increased running speed along the stem, but they also reveal a spatiotemporally separate ACx modulation of decision-making during maternal behavior. Although other auditory decision-making studies have seen ACx correlates of behavioral choice and action^46,47^, prognostic outcome modulation of ACx during learning has not been described to our knowledge. That this modulation was mainly suppressive was not explainable by task-engagement or movement^23–25^ since our animals were always engaged and moving similarly in both Correct and Wrong trials. Instead, our data were consistent with a lateral band suppression mechanism predicted from studies in passively listening mice^27,48^ but seen here in behaving animals, and likely provide a functional rationale for the observed importance of cortical inhibition in this reproductive behavior^14,42,44^. Speculatively, by helping to suppress potential distractor-related neural activity, prognostic suppression may substantiate an “attentional” mechanism for conveying a strong signal-to-noise message to downstream areas about a potentially behaviorally relevant stimulus^49^. Future studies would need to elucidate how selectively the suppression occurs for learned versus distractor stimuli.

While the magnitude of prognostic suppression increased over training, its strong prevalence on Day 1 was not expected. Afterall, the animal’s strategy for making a choice was not systematically altered until several days later. We hypothesize that the existing innate bias to follow a spatial-memory strategy in the absence of directional signals the animal would recognize as pup-related (e.g. USVs, odor, sight) must weaken first in order for a new sensory-cued strategy to take hold. We discovered a neural correlate of such weakening by decoding mPFC activity while animals returned to the nest – activity that was also predictive of the next-choice only in the first few days of training. Our results suggest a conceptual neural model (Figure 6j) for switching between different strategies. A brain region (or network) downstream of ACx and mPFC steers the animal either right or left at the T, and lacking sufficiently salient sensory information, persistently retains, while running back on Day 1 trials, a signal that determines the next-choice arm based on the spatial-memory strategy triggered by mPFC activity during retrieval. On some trials, the sound generates a strong enough signal-to-noise drive from ACx due to prognostic suppression that it updates this choice while the animal is running. Across days, this prognostic suppression along the stem strengthens, but also the initial mPFC seed representing the last-side information dissipates. Hence, a competition is created between the information conveyed by the two strategies. Identifying the brain area responsible for comparing these strategies remains as future work.

## Acknowledgement

This research was funded by National Institutes of Health grants R01DC008343 (RCL) and P50MH100023 (KL, RCL). We thank Li-Ling Shen for mouse colony maintenance; Hong Zhu and Dakshitha Anandakumar for their assistance with the chemogenetics experiment; and Malavika Murugan and Chris Rodgers for comments and insightful suggestions for the manuscript.

## Contributions

Research was conceived by RCL and KL. Experiments were carried out by KL, KW, LNZ, YTS and CJY. Data analysis was performed by KL and KW. Paper was written by KL and RCL, with feedback from all authors.

## Methods

### Animals

All mouse procedures were approved by the Emory University Institutional Animal Care and Use Committee. Animals were naïve female CBA/CaJ mice at age 8-10 weeks. Mice were socially housed in single-sex cages, on a 14h light / 10h dark reverse light cycle with ad libitum access to food and water.

### Chronic implant

In six mice, 16-channel silicon probes (A1x16-5mm-100-177-HZ16, NeuroNexus) were implanted through a craniotomy in the left ACx (Bregma 2.8 mm, ML 3.35 mm, 24 degree tilt, insertion depth 2.3 mm) under isoflurane (2%) anesthesia. Physiological status was monitored and maintained by a PhysioSuite (Kent Scientific) unit. In each surgery, two bone screws (000 X 3/32, J.I. Morris) were implanted above the cerebellum to be connected later to the ground and reference wire of the silicon probe. Silicon probes were inserted at a speed of 5 um/2s with a micromanipulator (M-10, Narishige). Then the craniotomy was covered with silicon elastomer (Kwik-Cast, WPI) then with dental cement (C&B Metabond, Parkell). In six other mice, silicon probes were implanted in the left mPFC (Bregma −1.7 mm, ML 0.7 mm, −10 degree tilt, insertion depth 2.5 mm) following the same procedures as for the ACx.

### T-maze setup

Behavioral training was performed in an 8’-2” X 10’-6” double wall anechoic chamber (IAC Acoustics) under dim red light. Mice were trained on an elevated T-maze ^1^ covered with Alpha-Dri (Shepherd Specialty Papers) bedding material. At the bottom of the stem, an 11 cm^2^ rectangular platform inset 1 cm lower than the stem served as a nest with extra bedding material. In each arm, a high frequency ribbon tweeter speaker (PT-R4, Pioneer) was placed 27 cm from the end of the arm and angled toward the intersection of the arms and stem. A third speaker (EMIT, Infinity) was positioned 30 cm above the nest with a slight tilt of 5 degrees toward the nest to mitigate standing wave interference. A heterodyne bat detector (Mini-3, Ultrasound Advice) was positioned 15 cm laterally away from the nest to record USVs on a RP2 processor (Tucker Davis Technologies) sampling at 20 kHz.

A camera (Flea 3, Teledyne FLIR) was positioned above the maze to record all behaviors (20 frames/s). The video recording was synchronized to the electrophysiology recording through a TTL trigger generated by the camera.

### Behavioral training

Prior to each training day, the T-maze was wiped down and clean Alpha-Dri bedding was added.

Before the implant surgery (Figure 2), animals were habituated to the T-maze for two days (30 min/day). On the next two days, two pups were positioned in the nest with the subject mouse during habituation (30 min/day). At the end of the habituation, most subject mice huddled on top of pups and thus were selected for chronic implants. Mice that did not huddle on pups or that cannibalized pups were excluded from further studies. Six days after recovery from the implant surgery, animals were re-habituated to the T-maze with two pups for an extra day (30 min).

At the start of each training day, two pups (postnatal P2 - P5 days, from a separate donor cage) were positioned at the nest so that the subject mouse would stay at the nest. Responses to pure tones presented from the overhead speaker at 60 dBSPL while animals remained at the nest were recorded to infer neural best frequencies. We presented 13 pure tones (4kHz −79kHz, 1/3 oct step, 0.12 s duration, 0.6 s inter-stimulus interval, with +/-0.15 s onset jittering, 30 repeats) in a pseudorandom order. Auditory stimuli were generated by Matlab (MathWorks) at the sampling rate of 223.214 kHz. Presentation of auditory stimuli were controlled by an RX6 processor running OpenEx (Tucker Davis Technologies).

As the training began, two pups were scattered on the T-maze to encourage the subject mouse to start searching^1^. The first sound pairing trial began the moment the subject mouse retrieved the last scattered pup back to the nest. The Target sound was an amplitude-modulated (5Hz) Gaussian noise, Butterworth bandpass filtered between 30-50 kHz. In each trial, the sound cue was presented from one of the two arm speakers. Once the subject mouse’s whole body (excluding the tail) crossed into the arm from the side where the sound was presented, a pup was delivered and the sound was terminated at the same time. If the mouse entered the opposite arm, then the sound presentation would continue until the mouse turned back and searched the arm playing the sound. Upon pup delivery, the mouse immediately retrieved the pup back to the nest, after which the next trial began.

The side where the sound and the pup would be delivered was predetermined pseudorandomly, so that the overall chance over each day’s 150 trials that each side was chosen was 50% and the probability that the same side was assigned consecutively in two trials was no more than 40%. The Target’s sound level was roved between 70 or 66 dB (calibrated at the T intersection) in a pseudorandom order with equal chance.

During electrophysiological recording, animals received 150 trials each day. In the chemogenetic experiment, animals received 100 trials each day to ensure all trials could be completed while CNO was active.

### Electrophysiology

A light tether cable was attached to the subject mouse’s implant through a ZF-16 zipf clip headstage (Tucker Davis Technologies) and thread through a commutator (ACO32, Tucker Davis Technologies). During passive listening and behavioral training, neural activities were recorded with a RX5 processor (Tucker Davis Technologies) at a sampling rate of 24,414Hz. Time series were band-pass filtered online between 100Hz −10kHz using the OpenEx environment.

### Histology

At the end of training and electrophysiological recording of each animal, two electrolytic lesions (10 uA, 15 s) were made at the first and last channel of the linear array. The animals were then sacrificed with CO2 and perfused with 0.1 m PBS and 4% paraformaldehyde. Coronal sections (50 um) were cut and stained with DAPI to localize electrode placement.

### Chemogenetics

We used AAV9-CaMKIIa-hM4D(Gi)-mCherry (titer: 1×10¹³ vg/mL, Addgene) and AAV9-CaMKIIa-EGFP (titer: 1×10¹³ vg/mL, Addgene) for chemogenetic silencing and viral control. hM4D(Gi) virus was infused into the ACx of 30 mice. EGFP virus was infused into the ACx of 8 mice. During viral injections, animals were anesthetized with isoflurane (2%). Craniotomies (1 mm diameter) were made above ACx of both hemispheres at the stereotaxic coordinate (ML +/-4.25, AP 2.80). A syringe (NanoFil, WPI) with a 36 GA needle was positioned at the center of the craniotomies. The needle was advanced into the cerebral tissue for 1.5 mm and was then backed up for 0.2 mm. Five minutes after needle insertion, virus (800 nL) was infused into the tissue at a rate of 100um/s (controlled by MICR 021, WPI). We then waited five minutes before withdrawing the syringe.

We waited three weeks before training animals with virus injections (Figure 1). The hM4D(Gi) + Clozapine N-oxide (CNO) group (N = 15 mice) was trained to test the effect of ACx silencing on learning.

Two others groups: hM4D(Gi) + Saline (N = 15) group and EGFP + CNO (N = 8) group were trained to serve as controls. Thirty minutes before training sessions began, 0.2 ml CNO (5 mg/kg, dissolved in DMSO, C0832 Sigma-Aldrich) or 0.2 ml saline was intraperitoneally injected into each mouse.

### Data analysis

#### Behavior tracking

Animal position and body parts were labeled automatically with DeepLabCut^2^ to extract frame-by-frame coordinates of animals in the maze.

#### Performance

An animal’s spatial-memory strategy usage was quantified as the percent of trials in which animals chose the side where it received the pup in the prior trial. Their sound-cued (auditory strategy) performance was quantified as the percent of trials in which animals chose the side where the sound was presented. An animal was considered to have learned the sound association once the percentage of trials combined over two days reached significance (p<0.05) by a binomial test.

#### Spike sorting

Single units were sorted offline with Kilosort2 and then manually verified and adjusted with Phy2^3^. Spikes that failed to be assigned to a single unit were classified into a multi-unit cluster on each channel. We did not assume the same units were recorded across days, though our statistical analyses (see below) did account hierarchically for units from the same animals.

#### Identifying sound-responsive ACx units

For each ACx unit, we compared its distribution of time-averaged spike rates during the sound ON to sound OFF periods. If a unit showed significantly different spike rates between these periods (criteria: p < 0.05, Wilcoxon signed-rank test), we considered the unit as sound-responsive.

#### Phase-locking power

The phase-locking power at 5 Hz was calculated from spike activities assigned in 1 ms bins during the sound ON period in a given trial. Neural activities were first smoothed in time with a gaussian filter with a 20 ms window, then the Welch’s power spectral density estimate was calculated with 1200 ms Hanning window (pwelch function in Matlab). For comparison, neural activities of the same trial were randomly shuffled across all time bins within the trial. Wilcoxon signed-rank test was used to compare the power calculated with the original responses versus the shuffled responses across the 150 trials of each recording session, with significant phase-locking when *p* < 0.05.

#### Best frequency (BF)

Because single unit responses to tones presented from above the animal were weak, best frequency was measured with multi-unit spike data at each recording channel and assigned to all units detected on that channel. First, single/multi-unit spikes recorded on each channel were combined. Second, auditory responses were quantified as the spike rate in the window from the tone onset to 30 ms after tone offset. The BF was defined as the frequency eliciting the highest response among all tones.

#### Spike rates in spatial bins

To measure the neural activities as an animal traversed the maze, spike rates were measured in time windows during which its center-of-head location fell into each 3.5 cm-long spatial bin along the path of traversal. Spike rates in spatial bins were calculated and analyzed separately during movement away from/back to the nest.

*Neural D’:* To quantify neural discriminability between two options, we computed a D’ for our various neural measures^4^. The equation for D’ is:

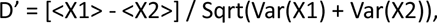

where X was 5 Hz phase-locking power (Figure 2) or firing rate (Figure 3 and 5). In most cases, discriminability was computed between Correct versus Wrong trials, though we also assessed D’ for Recorded versus Shuffled spikes (Figure 2b-e) and Left versus Right side choices (Figure 5b).

#### Identifying turn-selective mPFC units

To exclude mPFC units that were selective to turning, we computed the spike rates of left turn vs. right turn for each unit, on a trial-by-trial basis, when animals turn into the arms from the stem during the searching phase. Any units showing significant differences in spike rates between left vs. right turn (criteria: p < 0.05, Wilcoxon signed-rank test) were considered to be turn-selective units.

### Decoding

Neural activities in mPFC during retrieval were used to decode the pup location on the last trial or the side choice of the next trial. Turn-selective units were excluded from this analysis. First neural activities were calculated as firing rates as a function of location along the retrieval path, as described above. A decoder based on Long Short-Term Memory Networks^5^ (trainLSTM, Matlab, 10 hidden units, 20 epochs, batch size: 10) was used to classify trials based on spike rate vectors along the retrieval path. The decoding accuracy was evaluated by 5-fold cross-validation.

### Statistics

Because our neural dataset included the same animals, but not necessarily different units, recorded across days, we applied statistical analyses that could account for potential non-independence, as explained below. Statistical significance was set at p < 0.05.

#### Linear mixed-effects model

To determine effects of training at the level of individual units, we used a linear mixed-effects model (LMM) on all recorded units across all recorded animals from each day. In the model, each recorded Animal was treated as a random variable (ACx: 1-6; mPFC: 7-12) and each training Day (1-8) was treated as a fixed factor. A likelihood ratio test in MATLAB was used to compare the full model with the alternative model, in which the Day as a factor was removed.

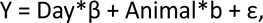

where Y was the 5 Hz phase-locking power or the firing rate or D’ for all units, depending on the application, β is the slope of the Day effect, b is the random effect coefficient, and ε is the error.

To determine effect of training on search speed at the level of trials, we also used a LMM with Day as a fixed factor and Animal as a random variable.

#### Linear regression

To determine effects of training at the session level, we averaged the activity across trials and units for each recording session, so that there were 8 values representing the 8 days for each animal. The 8 values were z-scored within each animal. Linear regression was performed with the 48 values (6 animals X 8 days) as a separate and more conservative test of whether there was an effect of training days.

To determine effect of training on search speed at the level of sessions, we used linear regression by averaging across trials for each recording session.

#### Hierarchical bootstrap

To determine whether there were overall significant effects in a measure for ACx or mPFC, irrespective of training day, we applied a hierarchical resampling and hypothesis testing protocol^6^. In each round of resampling, we first randomly selected with replacement one of the six subject animals, and then sampled with replacement from all recorded units of that selected subject animal. The resampling process was repeated 10,000 times. The population of bootstrapped means was then used to test hypotheses.

To test whether data from a single data set was significantly less (greater) than zero, we calculated the proportion of means of the 10,000 bootstrapped samples that were smaller (larger) than zero. If that proportion was greater than 95%, then the null hypothesis was ruled out.

To test whether the mean of one data set was significantly less (greater) than that of another, we obtained bootstrapped means for both datasets, and then calculated the proportion of means where dataset 1 was smaller (larger) than that of dataset 2. If that proportion was greater than 95%, then the null hypothesis was ruled out.

#### Generalized Linear Model

We tested the multifactorial influences on the firing rate of a given unit by constructing Generalized Linear Models^7^ with a log link function. We used

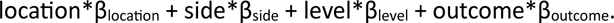

for the GLM predictors. These four factors were not correlated with each other, avoiding the problem of multicollinearity {Olsson}. A full model with all predictors was compared by a likelihood ratio test with four alternative models that were constructed by removing each factor from the full model. If the full model performed significantly better than the alternative model (p<0.05/(number of locations=22), for multiple comparisons correction), then the factor removed from the alternative model was considered as a significant effect in modulating the unit’s firing rate. To evaluate the false positive rate of our tests based on GLM models, we shuffled all trials and analyzed neural activities in the GLM models in the same way described above.

#### Other statistical tests

We used nonparametric tests whenever possible. The Spearman correlation was used to calculate correlation coefficients (r_s_). The Wilcoxon signed-rank test was used to compare two paired samples. The Wilcoxon rank sum test was used to compare two independent samples.

---

**Extended Data Figure S1.**
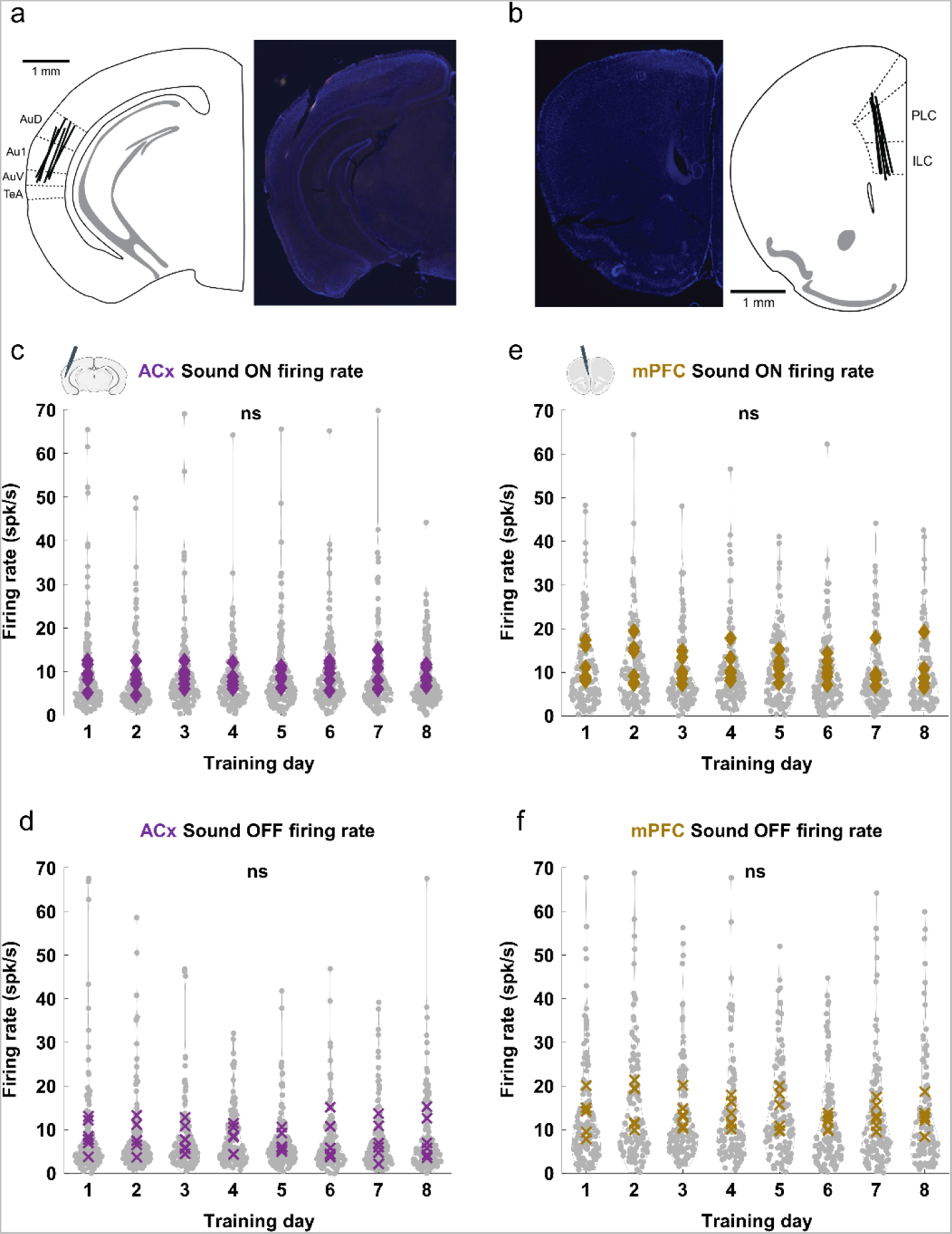
Firing rate of ACx and mPFC neurons during Sound ON and OFF did not change over days. a. Electrode tracks in ACx-implanted animals, plotted over a mouse atlas (Panel 54, {Paxinos}). AuD = Dorsal Auditory Cortex; Au1 = Primary Auditory Cortex; AuV = Ventral Auditory Cortex; TeA = Temporal Auditory Association Area. a. Spike rate of all ACx units during Sound ON showed no significant changes of the power across the 8 days training (linear mixed effect model LMM, LR = 0.516, *p* = 0.472). Each grey dot in the plot represents a unit. Each purple square indicates the mean of a recording session. b. Spike rate of all ACx units during Sound OFF showed no significant changes of the power across the 8 days training (LMM, LR = 1.562, *p* = 0.211). Each grey dot in the plot represents a unit. Each purple X indicates the mean of a recording session. c. Electrode tracks in mPFC-implanted animals, plotted over a mouse atlas (Panel 17, {Paxinos}). PLC = Prelimbic Cortex; ILC = Infralimbic Cortex. d. Spike rate of all mPFC units during Sound ON showed no significant changes of the power across the 8 days training (LMM, LR = 1.501, *p* = 0.221). Each grey dot in the plot represents a unit. Each brown square indicates the mean of a recording session. e. Spike rate of all mPFC units during Sound OFF showed no significant changes of the power across the 8 days training (LMM, LR = 2.867, *p* = 0.091). Each grey dot in the plot represents a unit. Each brown X indicates the mean of a recording session.

